# Phototactic Decision-Making by Micro-Algae

**DOI:** 10.1101/2025.07.02.662716

**Authors:** Shantanu Raikwar, Adham Al-Kassem, Nir S. Gov, Adriana I. Pesci, Raphaël Jeanneret, Raymond E. Goldstein

**Affiliations:** Laboratoire de Physique de l’École Normale Supérieure, ENS, Université PSL, CNRS, Sorbonne Université, Université Paris Cité, F-75005 Paris, France; Department of Chemical and Biological Physics, Weizmann Institute of Science, Rehovot 76100, Israel; Department of Physiology, Development and Neuroscience, University of Cambridge, Cambridge CB2 3DY, United Kingdom; Department of Applied Mathematics and Theoretical Physics, University of Cambridge, Wilberforce Road, Cambridge CB3 0WA, United Kingdom

## Abstract

We study how simple eukaryotic organisms make decisions in response to competing stimuli in the context of phototaxis by the unicellular alga *Chlamydomonas reinhardtii*. While negatively phototactic cells swim directly away from a collimated light beam, when presented with two beams of adjustable intersection angle and intensities, we find that cells swim in a direction given by an intensity-weighted average of the two light propagation vectors. This geometrical law is a fixed point of an adaptive model of phototaxis and minimizes the average light intensity falling on the anterior pole of the cell. At large angular separations, subpopulations of cells swim away from one source or the other, or along the direction of the geometrical law, with some cells stochastically switching between the three directions. This behavior is shown to arise from a population-level distribution of photoreceptor locations that breaks front-back symmetry of photoreception.

---

In areas as diverse as ecology [1, 2], microbiology [3], evolutionary biology and the psychology of human behavior [4] the question arises of how individuals make decisions when confronted with competing environmental stimuli. For complex organisms with a highly developed neural system, such decision-making may involve weighing the costs and benefits of the choices along with a balance between immediate rewards and long-term consequences. The situation is less clear for aneural organisms such as plants, bacteria and amoebae, but they appear to utilize similar mechanisms [5].

The simplest setting for decision-making clearly involves just two choices. At the scale of microorganisms there have been studies of the chemotactic response of the bacterium *E. coli* to two opposing chemical stimuli [3], and the phototactic response of colonies of cyanobacteria to two light sources [6–8]. These have suggested a “summation rule” in which the addition (scalar or vector) of the two stimuli forms the basis of the decision. Similar rules have been found in the study of plants [9]. Of the many types of taxes exhibited by microorganisms—chemotaxis, phototaxis, viscotaxis, durotaxis—phototaxis is distinguished by the fact that the stimulus direction and magnitude can be changed arbitrarily fast, without the complexities of a diffusive process or intervening surfaces. In the case of algal phototaxis, a long history of studies [10–22] has shown that the light-sensing process is “line of sight” in the sense that each cell has a photosensor that responds when directly illuminated, triggering changes in flagellar beating that produce alignment with the light. The two key ingredients for accurate phototaxis are the spinning of cells about a body-fixed axis and the directionality of the photoreceptor, achieved by a protein layer (the “eyespot”) that blocks light coming from behind the cell. Flagellar beating exhibits a rapid response to changes in light and a slower adaptation that is tuned to the spinning period [15, 16, 22, 23], and cells can exhibit positive or negative phototaxis depending on the prevailing light intensity.

Here we report on an extensive investigation at the single cell level of phototactic decision-making by unicellular green algae. By employing the experimental setup shown in Fig. 1(a,b), in which two collimated light beams with independently adjustable intensities intersect at a prescribed angle within a dilute suspension of negatively phototactic *Chlamydomonas reinhardtii*, we track thousands of individual cells’ decisions on the choice of swimming direction. The steady-state swimming directions are found to follow what we call the “tangent law”, an intensity-weighted average of the two light propagation vectors, and we show that this law is the fixed point of an adaptive model of phototaxis [19, 21, 22]. Studies of the response of cells to rapid changes in light direction reveal a surprisingly fast cell reorientation that can be quantitatively described as a limit of the adaptive theory. Finally, motivated by bifurcation phenomena found in certain decision-making processes [24], we examine swimming trajectories when the two lights are nearly antiparallel and find that there are three distinct subpopulations of cells: those that either (i) swim away from one source or from the other, (ii) go along the direction of the geometrical law, and (iii) exhibit stochastic switching between the first two choices. We show that this behavior is a consequence of a population-level distribution of the location of each cell’s photoreceptor relative to the equatorial plane of the cell.

**FIG. 1.**
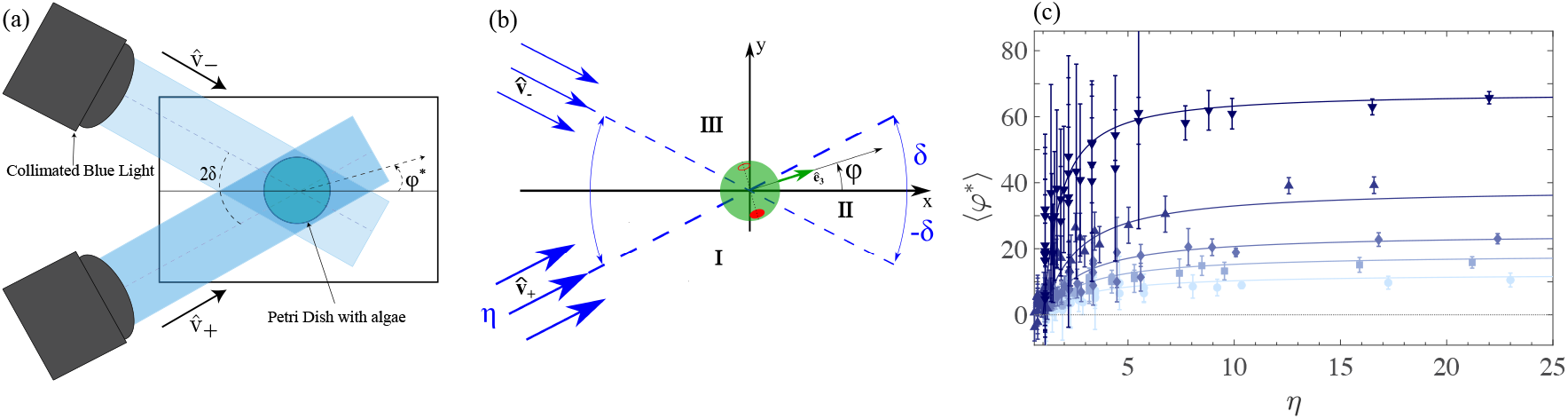
Experimental results. (a) Setup: two collimated lights shine toward the *x*-axis at angles ± *δ*, the lower with intensity *η* relative to the upper. (b) A swimming cell whose axis **ê**_3_ is at an angle *φ* with respect to the *x* − axis. Eyespot at two instants in time separated by a half cycle is shown as a solid and open red ellipse. (c) Swimming angle as a function of intensity ratio *η* for *δ* = 12.5^*°*^, 18.45^*°*^, 24.74^*°*^, 38.4^*°*^, and 67.6^*°*^ increasing upwards, along with the theoretical prediction (1) for each value of *δ*.

*C. reinhardtii* strain CC125 was grown axenically in Tris Acetate Phosphate (TAP) medium, which provides it with the required nutrients. Cells were grown at 22 ^*°*^C and synchronized in a light-dark cycle of 16*/*8 h (*∼* 70 *µ*E/m^2^s) with constant shaking at 160 rpm. They were harvested in the exponential phase (*∼* 10^6^ cells/mL) when they are the healthiest and most motile. The suspension was typically diluted by a factor of 20 to avoid collective effects, placed in an open Petri dish (Falcon 353001, diameter ∼ 3.5 cm) and kept in a dark box for 10 minutes before conducting experiments to ensure that all cells start from the same condition.

The experimental setup [25] consists of two collimated blue light beams (470nm, ThorLabs COP1-A) illuminating the Petri dish located at the center of the stage of an inverted microscope (Olympus, IX83), ensuring proper control of the light directions and avoiding light gradients over the imaging field (Fig. 1(a)). The light intensities were controlled by an LED Driver (Thorlabs DC4100) through their driving currents, and calibrated using a SpectraPen mini (Photon Systems Instruments). We use a 4 × objective (field of view 3.7 × 3.7mm^2^), and captured videos at 20 fps using a digital camera (Hamamatsu Orca Fusion-BT C15440-20UP). Image analysis used a combination of ImageJ and MATLAB to track the cells; trajectories were then linked and labeled employing the Crocker-Grier algorithm [26].

Because the algal suspension is contained in a thin chamber and the lights illuminate the chamber at the very shallow angle of ∼ 5^*°*^ with respect to the plane of the stage, the swimming is effectively two dimensional. As shown in Figs. 1(a,b), the two beams are at angles ±*δ* relative to the midplane, pointing toward the positive *x*-axis along the unit vectors 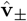, with *η* = *I*_+_*/I*_*−*_ *>* 1 the intensity ratio of the two beams.

*Chlamydomonas* cells, viewed from behind, spin counterclockwise around their posterior-anterior axis with frequency *f*_*r*_ = |*ω*_3_|*/*2*π* ∼ 1.5−2 Hz [25]. Because of shading by proteins behind the photoreceptor, when a cell swims at an angle *φ <* −*δ* (region I) or *φ > δ* (region III) both lights illuminate the photoreceptor in the same half turn, but when |*φ*| *< δ* (region II), the photoreceptor is illuminated only by one light in each half turn.

Starting from a dark state in which cells swim randomly, highly directional swimming occurs within ∼ 10 s of the start of illumination. The swimming paths appear as sinusoidal oscillations around linear motion because they are projections onto the *xy*-plane of helices. We obtain the swimming angle *φ*_*i*_ for each of typically 200 - 600 paths from the best-fit straight line and report ⟨*φ*^∗^⟩ as the ensemble average for each choice of (*δ, η*), with error bars representing the standard deviation of the fitted slopes. Figure 1(c) shows that the data are well-fit by the “tangent law”, where the swimming direction is along the unit vector **û**^∗^ = (cos *φ*^∗^, sin *φ*^∗^) defined as an intensity-weighted average of the light vectors,

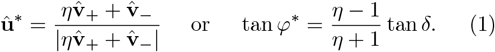

For *η* ≫ 1 (or *η* ≪ 1) the trajectory aligns with the lower (or upper) light (*φ*^∗^ = *δ* or *φ*^∗^ = − *δ*), while for equal light intensities (*η* = 1) cells swim along the *x*-axis (*φ*^∗^ = 0). This law appears to be valid for half-angles *δ* as large as ∼ 65^*°*^ −70^*°*^ (darkest blue inverted triangles in Fig. 1(c)), although in this situation cells do not follow the average direction as accurately, as illustrated by the increasing size of the error bars as *η* → 1 [25]. We hypothesize that such an intensity-weighted law should remain valid for more than two lights as long as the angle between the two furthest lights is small enough (2*δ* ≲ 140^*°*^).

While the result (1) makes no reference to biochemical processes in the cell, and is purely geometrical, we now show that it is a fixed point of a dynamical theory for *Chlamydomonas* phototaxis [22]. This theory combines rigid-body dynamics and an adaptive model for the angular rotation frequency *ω*_1_ around the body axis **ê**_1_ orthogonal to the flagellar beat plane due to asymmetries in beating of the two flagella in response to illumination of the photoreceptor. For a cell swimming in the *xy*- plane with a photoreceptor along **ê**_2_ in the cell’s equatorial plane (Fig. 4 in End Matter), and with *T* = |*ω*_3_| *t* a rescaled time, this dynamical system reduces to

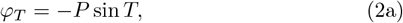

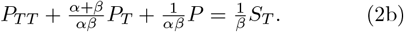

where the photoresponse variable *P* = *ω*_1_*/* |*ω*_3_|, and *α* and *β* are the slow flagellar adaption time and fast response time made dimensionless with |*ω*_3_|, respectively, with *α* ≫ *β* in experiments [22]. Under the assumption of additivity of light stimuli, the phototactic signal *S* = *P*^∗^ [*ηJ*_+_ℋ (*J*_+_) + *J*_*−*_H(*J*_*−*_)] is given by the projections 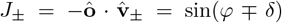 sin *T* of the two lights on **ô**, the outward normal to the eyespot, where 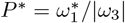, with 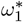 the peak turning rate around **ê**_1_. Here, the Heaviside functions ℋ in *S* represent the effect of eyespot shading. In the End Matter we show that averaging over the fast time scale of cellular spinning leads to the reorientation dynamics of the swimming angle *φ*,

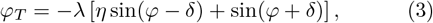

where *λ* ∝ *P*^∗^. While (3) depends on cellular parameters via *λ*, its steady state solution *φ*^∗^ is the tangent law (1).

The dynamics (3) has a Lyapunov function *V* (*φ*) in the sense that *λ*^*−*1^*dφ/dT* = −*dV/dφ*, with

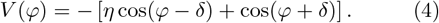

A simple calculation shows that 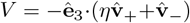, the projection of the total light vector onto the anterior pole of the cell, and that *V* is a minimum at *φ*^∗^; for negative phototaxis, a cell chooses a direction that minimizes the average light falling onto its anterior pole.

Moving on from the steady-state results in Fig. 1, we study how cells respond to a rapid switch in illumination between lights. Figure 2(a) shows cellular trajectories for 2*δ* = 70^*°*^. These turns are quantified through the angle *θ*(*T*) between the local unit tangent 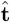 to the trajectory and the new light direction. Results for 5 different light intensities are shown in Fig. 2(b), which illustrates that complete reorientation occurs within just over one period of rotation (*T* ≃ 2*π*), with a modest dependence on light intensity. Within the adaptive theory, this rapid orientation corresponds to *P*^∗^ ∼ 1. When *α* ≫ *β* and *αβ* ∼ 1 as found in prior work [22], the dominant balance in (2b) gives *P* ≃ *S* and thus *θ*_*T*_ = *P*^∗^ sin *θ* H(sin *T*) sin^2^*T*. In terms of the shift 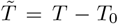 relative to the light-switching time *T*_0_, we find

**FIG. 2.**
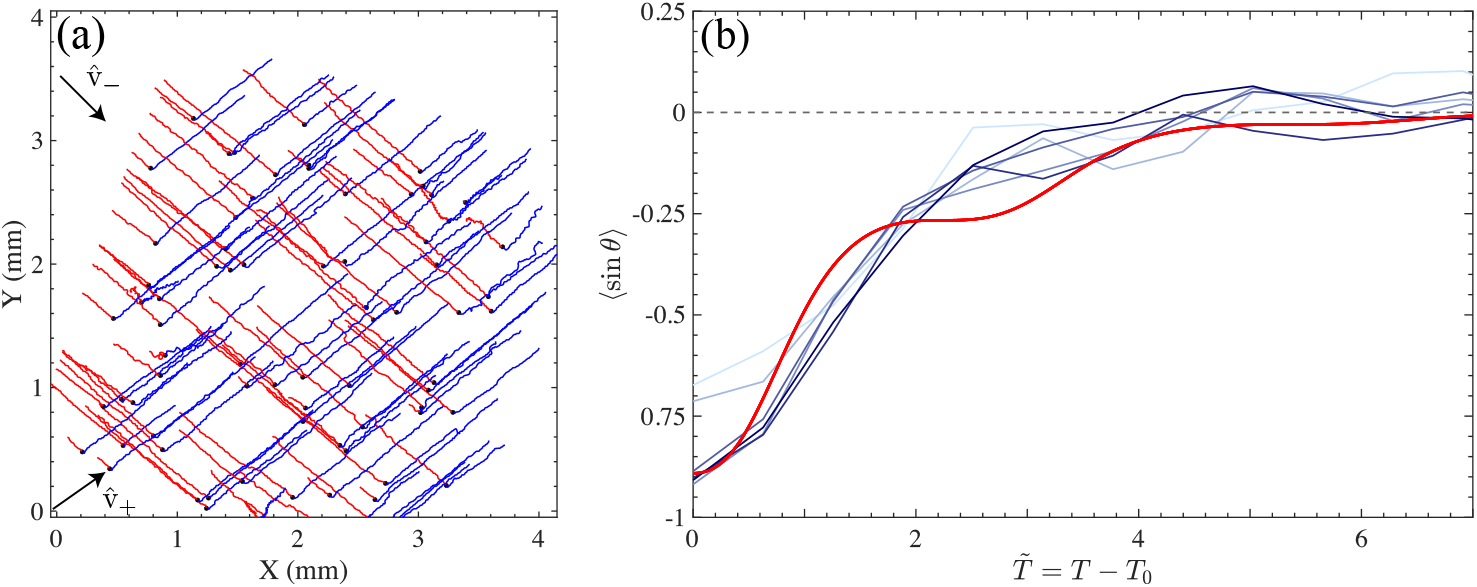
Reorientation dynamics. (a) Trajectories during a switch in light direction from 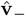 (red) to 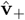 (blue) for *η* = 1. Black circle indicates time of switch. (b) Reorientation angle *θ* during phototurn at intensities *I* = 1.8, 3.7, 9.1, 17.7, 24.8, 33.9 W/m^2^, color-coded from light to dark blue compared to (5) (red) averaged over initial phase of motion.

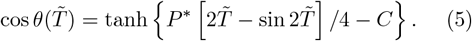

where *C* = ln(tan(*θ*_0_*/*2)) and *θ*_0_ is the initial angle. To compare with the experimental data, we note that when the lights are switched the cells are at random phases of their wiggly motion, and we average (5) over a uniformly distributed angle *θ*_0_ ∈ [− 110^*°*^ − *θ*_*i*_, − 110^*°*^ + *θ*_*i*_], where *θ*_*i*_ = 23^*°*^. The result shown in Fig. 2(b) matches the data well, with *P*^∗^ ≃ 1.4 as the sole fitting parameter.

This agreement validates the use of the simplified model for 2*δ* ≃ 180^*°*^ and *η* = 1, where rapid reorientations are relevant. The three types of trajectories described in the Introduction, shown in Fig. 3(a) for 2*δ* = 162^*°*^, are quantified in Fig. 3(b) by partitioning each into 3s segments (∼ 5 body rotations) and finding the average orientation angle *φ* for each. The probability distribution *Q*(*φ*) exhibits three peaks corresponding to swimming away from either light or along the *x*-axis.

**FIG. 3.**
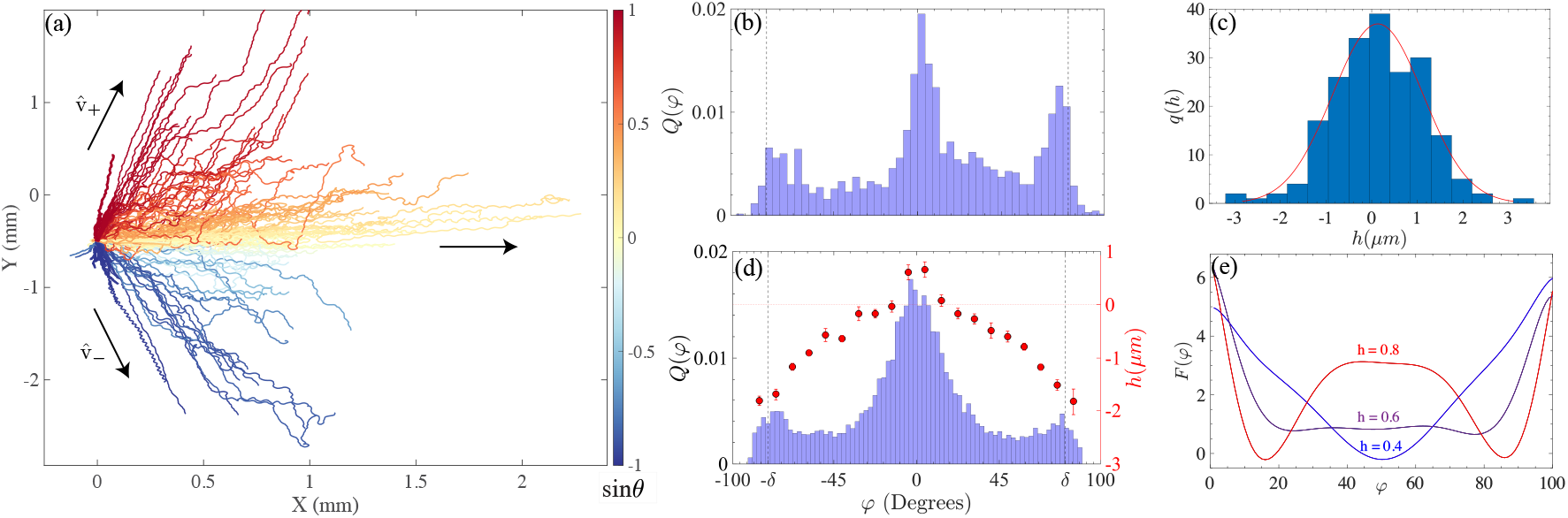
Phototaxis at large angular light separation. (a) Trajectories for 2*δ* = 162^*°*^ and *I*_*±*_ = 3.7W*/*m^2^, showing negative phototaxis, following tangent law, and stochastic switching between directions. Trajectories are color-coded by orientation of their end-to-end vectors relative to the *x* axis. (b) Probability distribution *Q*(*φ*) of trajectory angles for *I*_*±*_ = 3.7W*/*m^2^. (c) Distribution of eyespot offsets along with Gaussian fit. (d) Numerical *Q*(*φ*) from model of stochastic phototaxis incorporating distribution of eyespot shifts from equator. (e) Effective free energy as a function of eyespot offset for 2*δ* = 162^*°*^.

The theory discussed thus far cannot account for stable negatively phototactic swimming away from one light or the other when the photoreceptor is in the cell’s equatorial plane; the only stable fixed point is *φ*^∗^ = 0. But if the photoreceptor is displaced from the equator by an angle *γ <* 0, such motion is stable; due to photoreceptor shielding by the eyespot, a cell swimming away from one light must rotate through *γ* to sense the other. Such rotations may arise from flagellar beating jitter [22].

Using a method based on light reflection from the eye-spot [25, 27], we measured the probability distribution *q*(*h*) of eyespot displacements *h* from the midplane of 204 cells, and show in Fig. 3(c) that it is well-fit by a Gaussian with mean 0.15 *µ*m and standard deviation 1.0 *µ*m (positive values correspond to eyespots closer to the flagella). A significant subpopulation has eyespots displaced more than their own diameter from the midline.

It follows from the considerations above that during the finite duration of an experiment, cells with eyespots far below the midplane will have not had sufficient time to fluctuate enough to see the other light, and thus will swim away from one light or the other. For those with eye-spots far above the midline direct swimming away from either light is unstable. And those with intermediate positions can exhibit stochastic hopping between the three choices. To test the hypothesis that eyespot displacement is the origin of the distribution in Fig. 3(b) we generalize the model by writing a Langevin equation for the orientation with a shifted photoreceptor. The dynamical system is *φ*_*T*_ = − *P* sin *T* + ⟨*ξ*(*T*)⟩, where 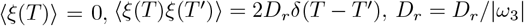 is the scaled rotational diffusion constant, the projections are

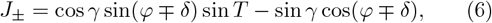

where *h* = *R* sin *γ*, with *R* = 5 *µ*m the cell radius, and we use the limit *P* ≃ *S* as in (5). In numerical studies we simulate *N* swimmers starting from random orientations in the plane, where each has an eyespot offset angle chosen from a Gaussian distribution with parameters taken from experiment. Figure 3(d) shows the resulting probability distribution function of trajectory angles, which strongly resembles the experimental one. We find that the peaks are associated with different subpopulations of the cells. Those with a large negative offset angle *γ* move directly away from one or the other light, while those with a strongly positive offset follow the tangent law. Cells with an offset close to the middle of these two extremes stochastically switch their trajectories.

By defining an effective free energy *F* = − ln *Q* from the measured distributions in the numerical computations we find an underlying bifurcation in the decision-making process. Simulations in which both *δ* and *h* are fixed show (Fig. 3(e)) a transition from a single minimum in *F* at small *h* to a double-well structure at larger *h*. Thus, the observed three-peak distribution function in Fig. 3(d), obtained for an ensemble of cells with different eyespot offsets, reflects a superposition of one- and two-minimum free energies. Further evidence for the existence of an underlying bifurcation at large angles is found in the increasing scale of fluctuations around the tangent law seen in Fig. 1(c) for large *δ* as *η* → 1 [25].

Since a continuously varying natural light field can be represented as a superposition of discrete sources, the results presented here suggest that the generalization of our model to one with *N* → ∞ lights would imply that negatively phototactic cells would swim away from the brightest spot. This result provides a linkage between the line-of-sight and gradient-climbing approaches [28–30]. Issues for further study involve the possibility of a dynamic bifurcation [31] when the lights are point sources, not at infinity, and the apparent position slowly varies as cells swim [24], as well as the effects of longer-term adaptations associated with photosynthesis [32, 33].

We thank Nelson Pesci for discussions. This work was supported in part by ANR JCJC funding (ANR-23-CE30-0009-01; RJ & SR), the Lee and William Abramowitz Professorial Chair of Biophysics and a Royal Society Wolfson Visiting Fellowship (NSG), the John Templeton Foundation and the Complex Systems Fund at the University of Cambridge (AIP & REG).

## Data Availability

The data that support the findings of this article are openly available [34].

## END MATTER

The complete model introduced earlier [22] consists of the coupled dynamics of the Euler angles (*ϕ, θ, ψ*) describing rigid body motion and the adaptive dynamics of the angular rotational velocity *ω*_1_ about the axis **ê** _1_ that is orthogonal to the plane of the flagellar beating. In a system of units scaled by the magnitude of the spinning angular velocity |*ω*_3_| about the cellular axis **ê** _3_ that lies in the plane of beating and is equidistant to the bases of the two flagella, the dynamics takes the form

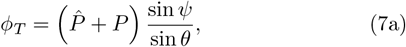

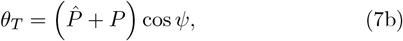

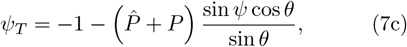

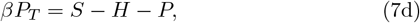

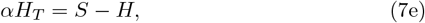

where 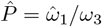 and *P* = *ω*_1_*/ω*_3_ are the scaled intrinsic and photoresponse rotation frequencies around **ê**_1_, respectively, *H* is the hidden variable responsible for adaptation and *α* and *β* are the scaled response and adaptation times for the photoresponse system.

In the absence of any light signal (*S* = 0) the photoresponse variables vanish, we can describe analytically a weakly helical trajectory associated with a small 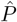 that deviates weakly from the *x* − *y* plane. Setting 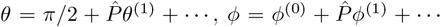, and observe that 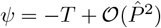. A simple calculation yields *ϕ*^(1)^ = cos *T* and *θ*^(1)^ = sin *T*, thus

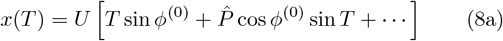

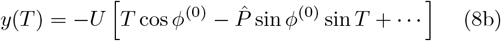

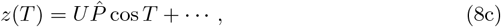

the equation of a helix about a line in the *x* − *y* plane oriented at angle *ϕ*^(0)^ with respect to the *y*-axis. If we define **û** = (sin *ϕ*^(0)^, −cos *ϕ*^(0)^) and **û** ^⊺^ = (cos *ϕ*^(0)^, sin *ϕ*^(0)^), then the position of the swimmer is

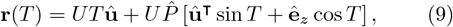

making clear that the helical radius is set by 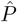. As this solution holds when *S* = 0, we can view it as the solution to the “homogeneous” linearized version of Eqs. (7). Hence, the full solution of the linearized inhomogeneous problem is the sum of (8) and the particular solution of the linearized inhomogeneous problem.

In earlier work [22] we found two intrinsic time scales of relevance to a phototactic turn: a short one associated the cell’s spinning motion about the axis **ê**_3_ and a longer one for the reorientation of **ê**_3_ toward a collimated light beam. In this section, we provide a systematic derivation of the averaged dynamics based on the presumed smallness of the photoresponse variable *P*, focusing first on the case of a single light source pointing along −**ê**_*x*_. To make analytical progress, we ignore the small helical component of the motion, take the photoreceptor to be along **ê**_2_, and fix the Euler angle *θ* = *π/*2 to neglect the small out-of-plane motion. Finally, with *P* small, we may approximate the third Euler angle by *ψ* = −*T* since the time evolution of *ψ* is dominated by rotations around **ê**_3_. This yields the simplified model for the Euler angle *ϕ*

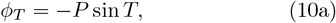

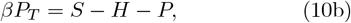

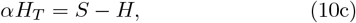

We set *P*^∗^, the scaled maximum magnitude of the photoresponse, to be

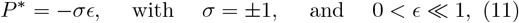

±1 for positive/negative phototaxis. Then the signal is

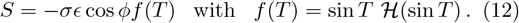

Noting that (10b) and (10c) can be combined into a single second-order equation for *P* [22],

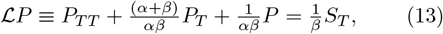

we now seek a perturbative solution to the dynamics (10a) and (13) in powers of *ϵ*.

As the intrinsic rate of variation of *P* given by *α* and *β*, is 𝒪 (1) it is natural to assume that when coupled to the slow variable *ϕ* the dynamical variables *ϕ* and *P* can be written as functions of two variables, a fast one (*T*) and a slow one (*τ* = *ϵT*). Thus assuming the forms *ϕ*(*T, τ*), etc. we have the rule

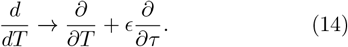

If we now propose perturbative expansions of the form

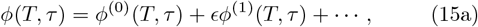

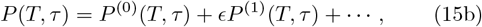

then at 𝒪 (*ϵ*^0^) we find

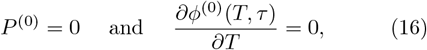

which implies that *ϕ*^(0)^ depends only on the slow variable *τ*. This solution corresponds to the fixed point of the underlying adaptive dynamics in the absence of a stimulus.

At 𝒪 (*ϵ*^1^) we find

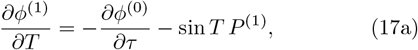

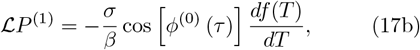

Note that the r.h.s. of (17a) does not depend on *ϕ*^(1)^ and thus functions as a forcing term. Moreover, at this order the right-hand-side of (17b) is a functios of *T*, with *τ* playing the role of a parameter. It is thus possible to solve (17b) independently of (17a). The l.h.s. of (13) is simply a damped harmonic oscillator. Since [(*α* + *β*)*/*2*αβ*]^2^ *>* 1*/αβ*, the Green’s function is

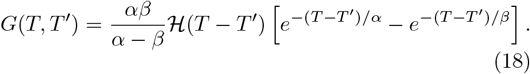

The term *df /dT* on the r.h.s. of (17b) is cos *T* (sin *T*) + cos *T* [sin *T δ*(sin *T*)], whose second term makes no contribution to the convolution with *G*. With initial condition *P* ^(1)^(0) = 0, the solution at a time *T* such that 2*nπ < T* ⩽ 2(*n* + 1)*π* (i.e. *n* − 1 full cycles have passed) is 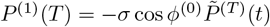, where

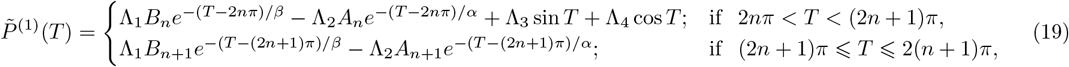

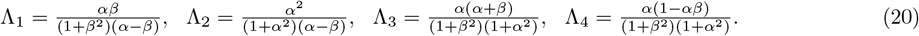

For the realistic parameters *α* = 7 and *β* = 0.14 [22], Fig. 5 shows the function 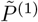 over several complete cycles.

**FIG. 4.**
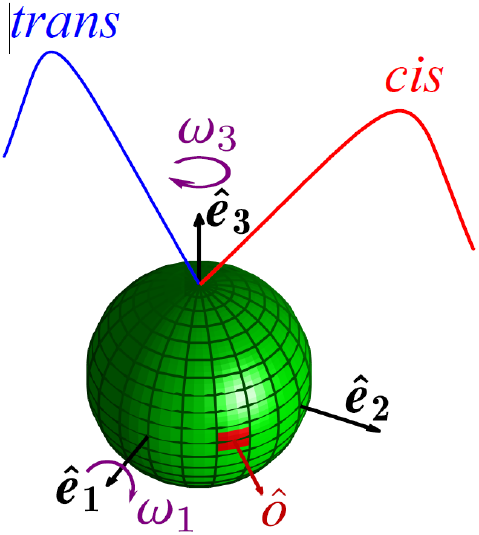
Coordinate system of *Chlamydomonas*.

**FIG. 5.**
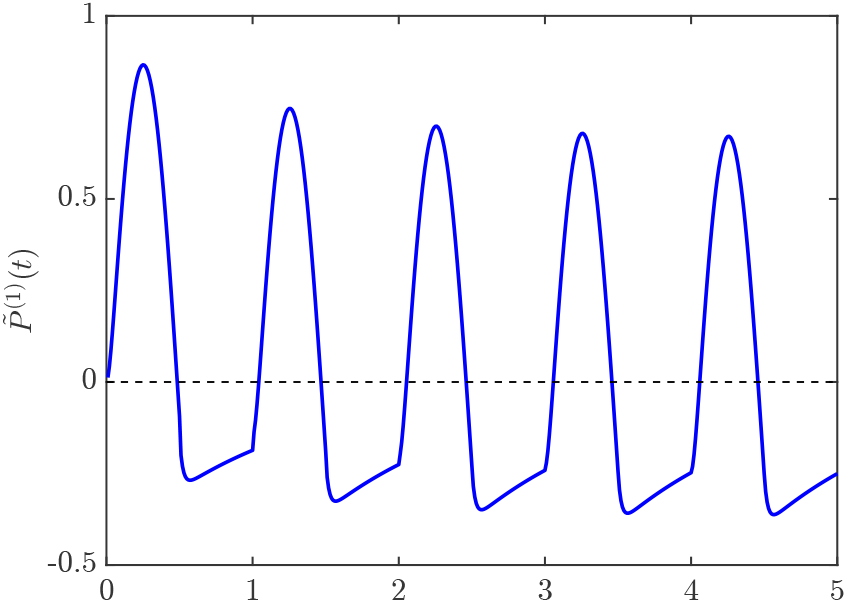
The correction function 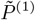 (1) for α = 7 and β = 0.14.

Since we know that the physically relevant solution for the angle *ϕ* is bounded as *T* →∞, we must seek a bounded solution to (17a), which can only be achieved by removing any secular terms. The freedom to do so is provided by the as yet unknown value of *∂ϕ*^(0)^*/∂τ*.

The solvability condition is that the r.h.s. of (17a) be orthogonal to the nullspace of the l.h.s. operator. As the nullspace is a constant, we require only that the vanishing of the integral of the r.h.s. over any complete period from *T* = 2*mπ* to 2(*m* + 1)*π*, for *m* = 0, 1, The necessary integrals were performed in earlier work [22], from which we deduce the equation of motion necessary for a bounded solution,

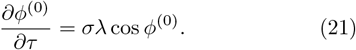

where λ = Λ3/4. Expressed in terms of the angle φ^(0)^ = ϕ^(0)^ − π/2, with initial condition φ_0_ = φ^(0)^(0), we have

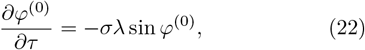

whose solution is

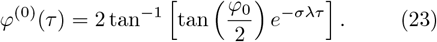

For positive phototaxis (*σ* = +1), *φ* → 0 as *τ* → ∞ (the cell swims along +**ê**_*x*_, antiparallel to the light) whereas *φ* → *π* as *τ*→∞ for negative phototaxis (*σ* = − 1), as the cell swims parallel to the beam.

The generalization of (22) to the case of two lights at angles ±*δ*, leading to (3), is straightforward.

## SUPPLEMENTAL MATERIAL

### S1. FOURIER ANALYSIS OF CELL TRAJECTORIES IN SINGLE-LIGHT EXPERIMENTS

To understand in detail the effect of light intensity on the swimming of the cells, we first studied the trajectories in response to a single collimated light beam. Experiments were performed using a × 4 objective at 90 fps in order to obtain well resolved Fourier spectra of the motion. Because cell trajectories present modulations at frequencies below 0.5 Hz when cells follow a single light direction (Fig. S1(a), solid blue line), we first applied a Savitzy-Golay (SG) filter to the tracks in order to define properly the underyling mean motion around which are fluctuations associated with the the helical motion of the cells. The parameters of the SG filter were fixed to window size of 49 frames and polynomial order as 1. The SG filter then yields a mean trajectory (dashed black line in Fig. S1(a)) around which we can analyze the fluctuations of the filtered path, as shown in Fig. S1(b). The Fourier transforms for several light intensities shown in Figs. S1(c,d) are obtained by first trimming all trajectories at a given intensity to a common duration of 1300 frames, computing the FFTs of each, and then averaging those spectra. We observe a clear peak at 1.5 − 2 Hz, common to all intensities, that corresponds to the frequency of helical swimming. Interestingly, the structure of the spectra above 2 Hz appears to depend on the light intensity; at low intensity there is a small peak (or “shoulder”) between 3.5 and 4 Hz that disappears as the light intensity is increased, while other peaks appear at just below 6 and 8 Hz. Other peaks are also present at larger frequencies (between 10 and 20 Hz) as shown in Fig. S1(c). The biological explanation for these high-frequency peaks and their dependence on light intensity remains unclear. The broad peaks seen between 30 and 40 Hz, corresponding to the small, fast fluctuations visible in Fig. S1(d), arise from detection noise of the positions of cells.

### S2. TANGENT LAW

#### A. Current-to-Power conversion

The LED driver controls the electrical current sent to the device. To calibrate the power transmitted by the LED, we used a power meter (Thorlabs controller PM100D with S130VC power sensor) that is held perpendicular to the collimated light beam, at the center of the stage, where we image the algae. We perform a linear fit to the plot of power versus current, the slope of which yields the calibration coefficient that is used in converting the ratio *i*_+_*/i*_*−*_ of the electrical currents to a ratio of light power or light intensity *η* = *P*_+_*/P*_*−*_ = *I*_+_*/I*_*−*_. This measurement is repeated over all experiments with different angles. An example of a calibration curve is shown in Fig. S2.

**FIG. S1.**
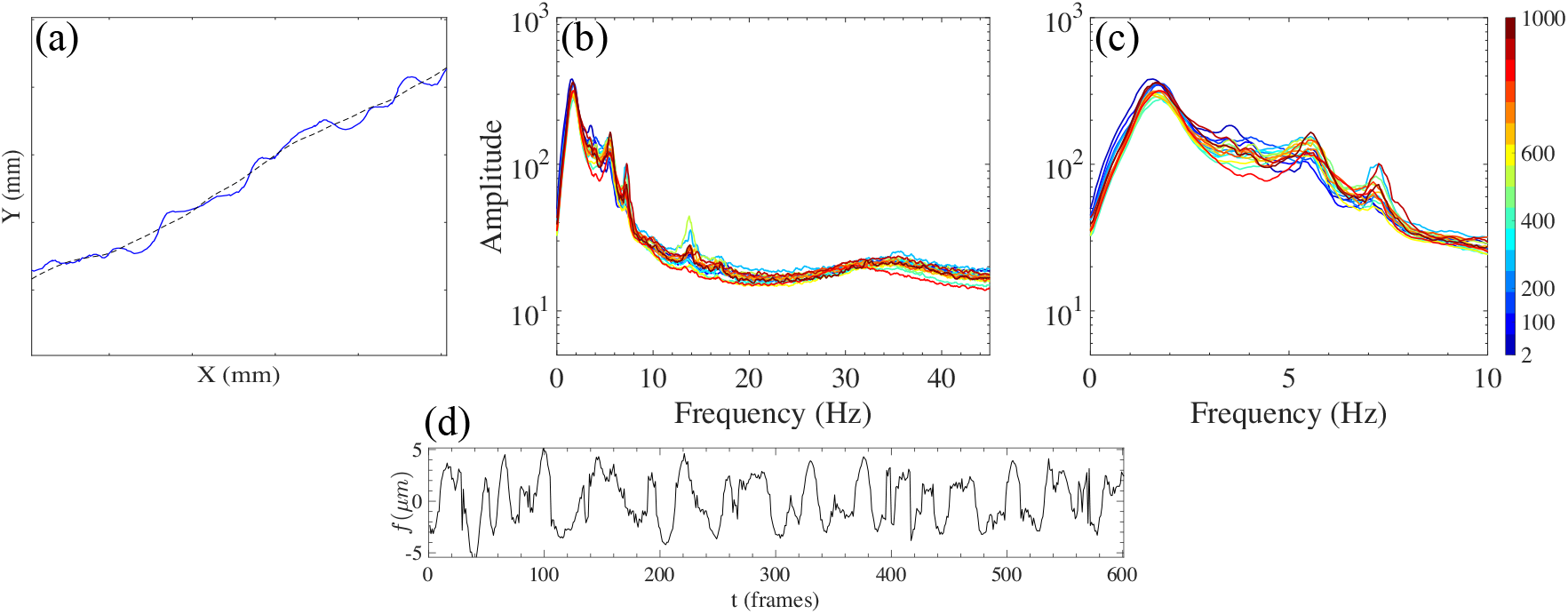
Analysis of swimming trajectories. (a) Typical alga trajectory (solid blue) with its mean computed from the SG Filter (dotted black). (b) Fourier spectra for different light intensities (c)Enlargement of the first 10 Hz of the spectra shown in (c) (see colorbar). (d)Fluctuations of the trajectory shown in (a) around the mean trajectory computed from the SG filter.

**FIG. S2.**
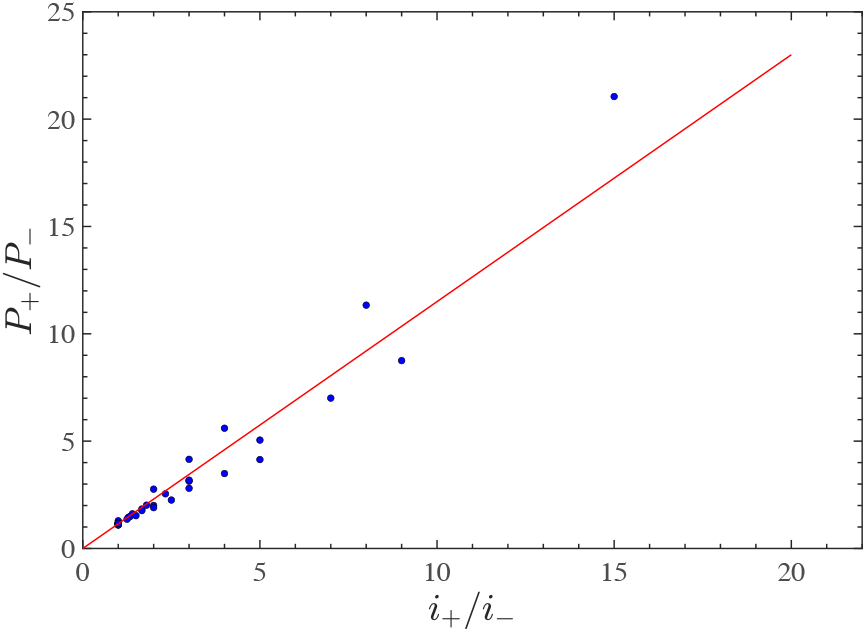
Calibration curve for 2*δ* = 49.5^*°*^ with the linear fit shown in red (here *P*_+_*/P*_*−*_ = 1.15 *i*_+_*/i*_*−*_).

**FIG. S3.**
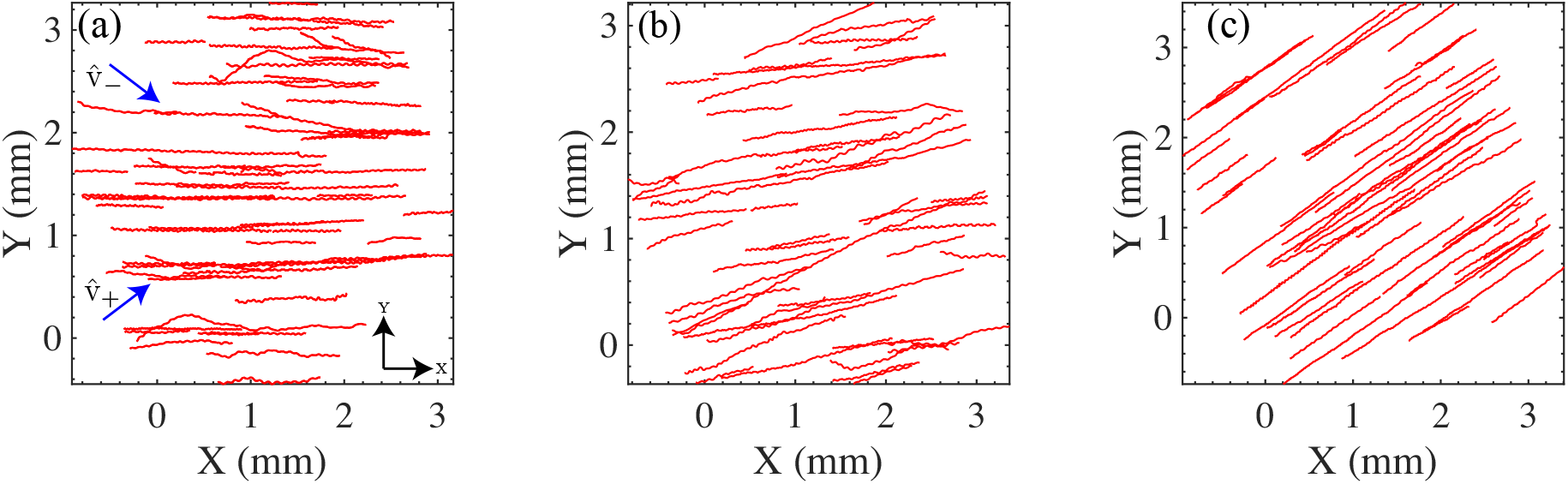
Trajectories for 2*δ* = 76.8^*°*^ under different light intensity ratios: (a) *η* = 0.74, (b) *η* = 2.9, (c) *η* = 16.6.

For information, with our blue LED sources the conversion of light intensity in W*/*m^2^ to *µ*E*/*m^2^*/*s is given by: 1W*/*m^2^ = 3.96*µ*E*/*m^2^*/*s.

#### B. Experimental determination of the angles for the tangent law

To obtain the tangent law, two series of experiments were performed for each fixed position of the two lights. We first performed tracking experiments with each light turned on independently, which allows us to get an accurate value of the angle 2*δ* between the lights in the frame of the camera. This relies on the fact that cells do follow accurately the light propagation direction when a single light source is on. The ensemble average of the swimming direction of many cells, each obtained by a linear fit to its trajectory, is then used to define the light propagation directions. We then performed experiments with the two lights turned on to extract the ensemble average angle of swimming ⟨*φ*^∗^⟩ as a function of *η*, obtained also by a linear fit to the trajectories. Figures S3(a-c) show representative trajectories of the cells for a fixed 2*δ* = 76.8^*°*^ at different values of *η*, illustrating how the swimming angle *φ*^∗^ changes as *η* increases.

#### C. Analysis of the standard deviation of *φ*^∗^

As the angle between the two lights is increased to 2*δ* ≈135^*°*^, we observed that the fluctuations in cell swimming direction drastically increased as *η* gets closer to 1 (Fig. S4). Although the tangent law can here still be considered as valid on average, the ability of cells to follow the intensity-weighted average light direction is impeded at such large angles. This is a first sign that a bifurcation will be observed for larger angles, as we see for 2*δ* = 162^*°*^. As discussed in the main text, the three-peak distribution of trajectory angles arises from a superposition of the properties of subpopulations of cells. For this reason, there should be no sharp transition or true bifurcation as *δ* is increased.

**FIG. S4.**
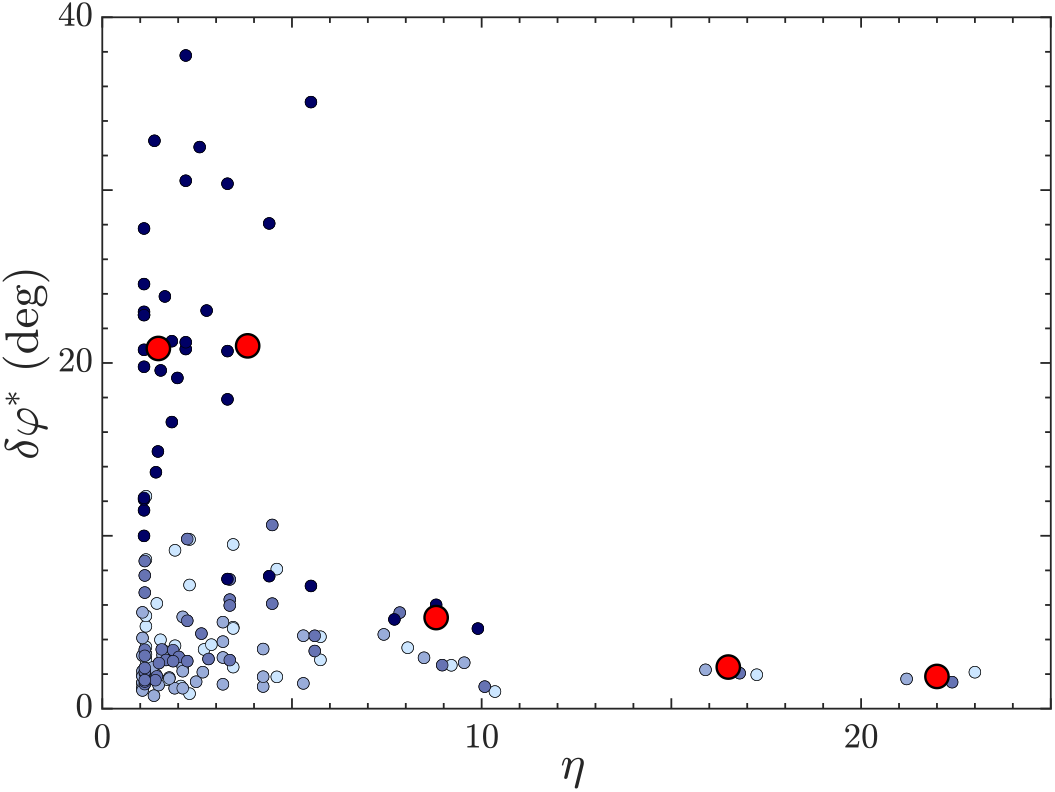
Standard deviation of swimming direction *δφ*^∗^ among the cell population for all *δ* (color-coded from light to dark blue for increasing *δ*, as Fig. 1(c) of main text). For the largest angle 2*δ ≈* 135^*°*^ (dark blue) the standard deviation increases drastically as *η* → 1 (for *η* ≲ 5). This is shown by the red circles, obtained by binning the data with similar values of *η*.

### S3. LIGHT-SWITCHING EXPERIMENTS

#### A. Obtaining the switch time *t*_0_

The experiments were performed at a frame rate of 20 fps, using the same 4× objective as for experiments above. Movies were recorded for a period of 20 seconds, within which the light was switched manually at ∼ 10s after the start of the recording. To determine the video frame corresponding to the switching of the light, and hence the switch time *t*_0_, we located the jump in total intensity of the video frames that arises from the very small intensity difference of the two light sources. Fig. S5 shows a typical intensity trace with such a jump.

#### B. Dispersion of initial value of sin *θ* in light-switching experiments

In the light-switching experiments, the angle of swimming of the cells at the switch time *t*_0_ does not correspond exactly to the light propagation direction 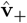. This is a consequence of the helical motion of the cells that makes their orientation oscillate around the average direction 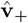. Typical distributions of orientations before the switch are shown in Fig. S6 for *I*_+_ = 1.8W*/*m^2^ (*i*_+_ = 5mA) (panel a) and *I*_+_ = 63.1W*/*m^2^ (*i*_+_ = 100mA) (panel b). In order to compare with the theory (Fig. 2b and Eq. (5) of the main text), we need to take into account this variation in orientation at *t*_0_, which makes the average value ⟨sin *θ*(*T*_0_)⟩ larger than − sin 2*δ* ≈− 0.94. The width of these distributions decreases as the light intensity increases (Fig. S6), consistent with the ordering of the curves in Fig. 2(b) of the main text at the switch time *t*_0_.

**FIG. S5.**
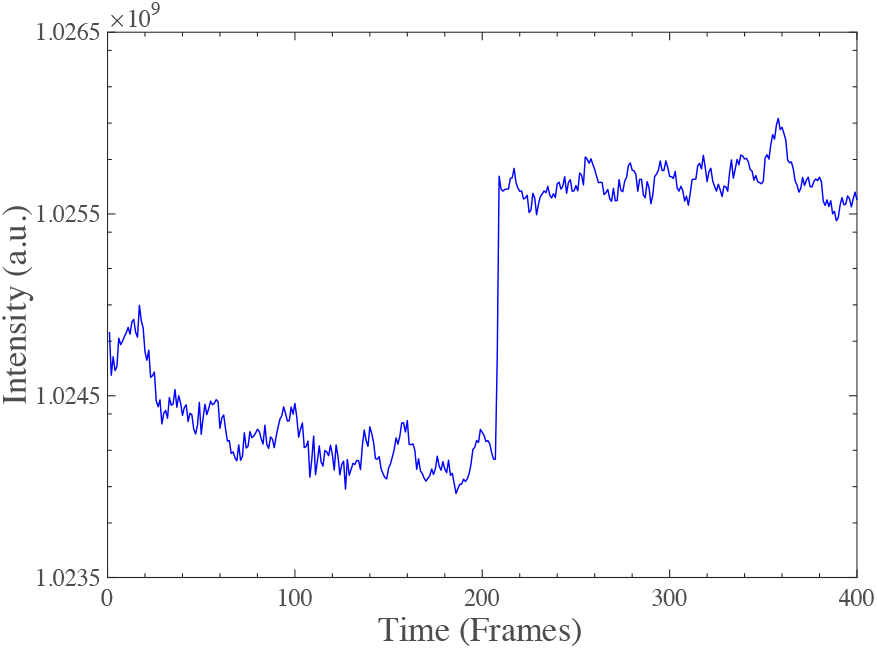
Total frame intensity over the duration of an experiment for *i*_+_ = *i*_*−*_ = 50 mA.

**FIG. S6.**
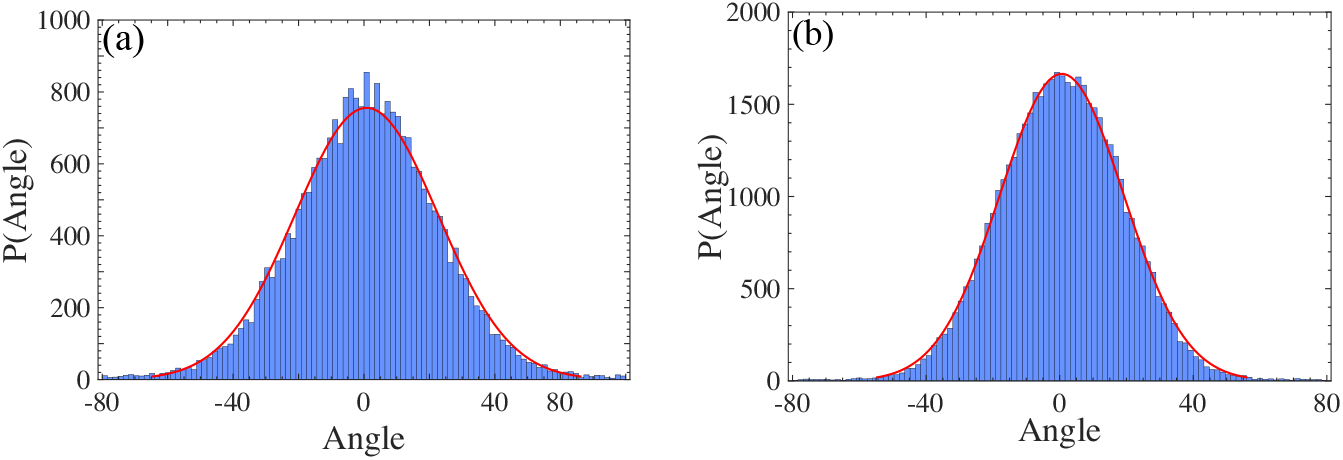
Distribution of swimming directions prior to the switch time *t*_0_ in light-switching experiments. The distributions are centered around the direction of light propagation 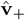. (a) A Gaussian fit (red curve) at low intensity *I*_+_ = 1.8W*/*m^2^ (*i*_+_ = 5mA) yields the average angle *µ* = 0.78^*°*^ and standard deviation *σ* = 21.89^*°*^. (b) At the larger intensity *I*_+_ = 63.1W*/*m^2^ (*i*_+_ = 100mA), *µ* = 0.63^*°*^ and *σ* = 18.52^*°*^.

### S4. BIFURCATION

#### A. Calibrating light intensity

For the bifurcation experiments at large angles 2*δ* between the lights, we require the intensity of each light to be exactly the same. Otherwise, there would be a strong asymmetry in the distribution of orientations *Q*(*φ*) because cells are highly sensitive to even minute differences between the two light intensities.

To ensure that *η* is as close to 1 as possible, we calibrated the two LEDs using a Spectra Pen Mini (Photon System Instruments). A custom stage was built to hold the sensor at its center, exactly where the cells are imaged in the Petri dish. We then adjusted the electrical current sent to one of the lights to match exactly the intensity of the second light. This protocol leads to distributions *Q*(*φ*) that are quite symmetric, although a slight asymmetry persists.

#### B. Calculating the distribution of swimming directions *Q*(*φ*)

To obtain a detailed quantification of the swimming direction of the cells, we divide each trajectory into segments consisting of 5 full rotation periods (60 frames = 3s). From these segments we calculate the local angle (*φ*) with respect to the average direction 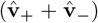, yielding the distribution of orientations (*Q*(*φ*), Fig. 3(b) in the main text). The same analysis was performed on the simulated trajectories (Fig. 3(d) in the main text).

#### C. Eyespot imaging

In order to determine the distribution of eyespot locations in a population of *C. reinhardtii*, we first immobilized the cells on a microscope slide coated with polylysine (Electron Microscopy Sciences, #63410-01) by placing a 12 *µ*L droplet onto the slide and covering it with a coverslip. We then combined the information obtained from three different types of imaging with a 40× objective (Olympus, UPlanSApo, 40× /0.95): brightfield microscopy, reflection microscopy, and epi-fluorescence microscopy (methods detailed below). Brightfield imaging (Fig. S7(a)) was used to characterize the cell shape. Reflection microscopy with an orange (580nm) light (Fig. S7(b)) was used to image the eyespot as a bright spot making use of the highly reflective properties of the carotenoid layer of the eyespot at this wavelength. Epi-fluorescence microscopy was finally used to image the chloroplast and obtain the polarity of the cells. Acquiring about 20 images with these methods, we obtained statistics of the eyespot location of ∼ 200 cells. A superposition of the three types of images is shown Fig. S7(f) with the brightfield image in grey, the eyespot in red and the chloroplast in green.

**FIG. S7.**
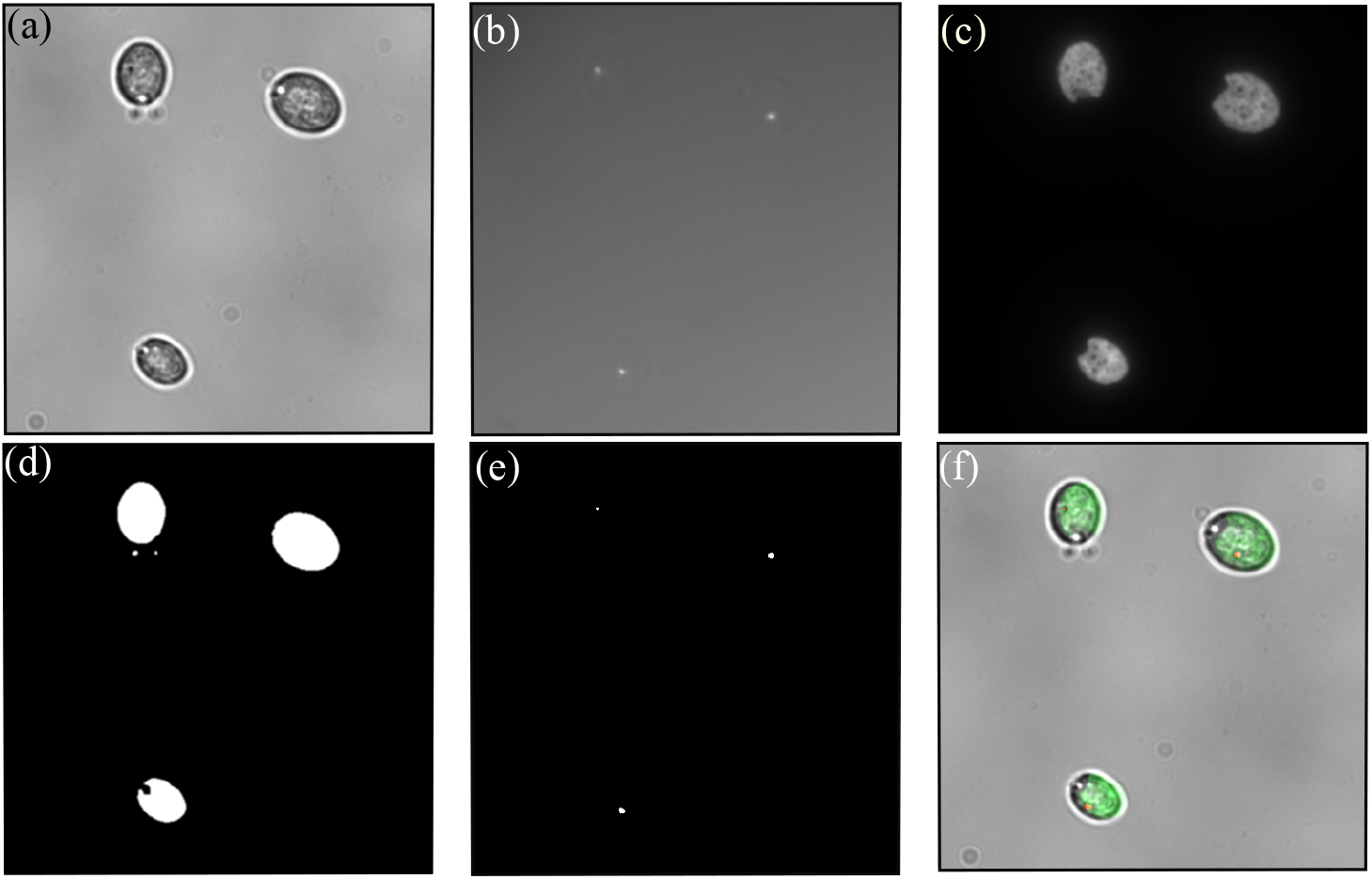
Determining the eyespot location. (a) Brightfield image of the cell bodies, (b) eyespots (bright spots) in reflection microscopy, (c) chloroplast visualized using epi-fluorescence microscopy of the chlorophyll molecules, (d) binarized image of panel a, (e) binarized image of panel b, and (f) merged image of panels a, e and c in gray, red and green color respectively.

The brightfield images were binarized, followed by dilation and erosion to obtain ellipsoidal cell bodies (Fig. S7(d)) from which we extracted the centroid, the major and minor axis, the cell area (to filter out cells too close to each other to be detected as single cells) and the bounding rectangle of the features. The images from reflection microscopy were also simply binarized (Fig. S7(e)) to extract the eyespot position. From the information of the eyespot position and the major axis of the cells we obtained the distance of the eyespot from the cell equator, without knowledge of the direction (i.e. closer to the flagella or further away). This missing information was obtained from the epi-fluorescence images of the chloroplast which makes a U-shape at the back of the cells (Fig. S7(c)). Extracting the center of mass of the fluorescence signal allowed us to know the polarity of the cells. To do so we applied the binarized brightfield images as masks to the epi-fluorescence images, then cropped around each cell (using the bounding rectangle) and finally computed the center of mass of the fluorescence signal.

#### Details of the imaging methods

All images were acquired on an Olympus IX83 inverted microscope equipped with a Hamamatsu Orca Fusion-BT camera (C15440-20UP).

*Brightfield*. Brightfield images were captured using a red filter at 624 nm (BrightLine, FF01-624/40-25) to ensure cells remained still by avoiding phototactic responses.

*Reflection microscopy*. White light from a Lumencor SOLA-VN LED source was sent through the back of the microscope into a filter cube containing an excitation filter at 580nm (BrightLine, FF01-580/60-25) and a glass slide acting as a semi-reflective surface to redirect light towards the sample and let the reflected light from the eyespots to reach the camera.

*Epi-fluorescence microscopy*. White light from a Lumencor SOLA-VN LED source was sent through the back of the microscope into a filter cube containing an excitation filter at 447nm (BrightLine, FF02-447/60-25), a dichroic mirror at 605nm (Thorlabs, DMLP605R) and an emission filter at 697nm (BrightLine, FF01-697/58-25) to collect the fluorescence signal of the chlorophyll molecules in the chloroplast.

### S5. CAPTIONS FOR SUPPLEMENTARY MOVIES

**Supplementary Movie 1:** Cells following the tangent law, at various values of *η* = 1.2, 2.9, 16.6. Video captured at 4x magnification, 20fps, played in real time.

**Supplementary Movie 2:** On changing the light, we see the fast turns that the cells make when the light is switched from 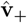 to 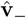. Video captured at 4x magnification, 20fps, played in real time.

**Supplementary Movie 3:** At bigger angle of 2*δ* = 162^*°*^ and *η* = 1 different subpopulations of cells emerge: cells following either light, cells following the tangent law and cells switching stochastically between these two. Video captured at 4x magnification, 20fps, played in real time.

## Notes

https://doi.org/10.5281/zenodo.15051634

### Competing Interest Statement

The authors have declared no competing interest.

